# Nanotopography enhances dynamic remodeling of tight junction proteins through cytosolic complexes

**DOI:** 10.1101/858118

**Authors:** Xiao Huang, Xiaoyu Shi, Mollie Eva Hansen, Cameron L. Nemeth, Anna Ceili, Bo Huang, Theodora Mauro, Michael Koval, Tejal A. Desai

## Abstract

The epithelial tight junction regulates barrier function and is responsive to extracellular stimuli. Here we demonstrated that contact of synthetic surfaces with defined nanotopography at the apical surface of epithelial monolayers increased paracellular permeability of macromolecules. To monitor changes in tight junction morphology in live cells, we fluorescently tagged the scaffold protein zonula occludens-1 (ZO-1) through CRISPR/Cas9-based gene editing. Contact between cells and nanostructured surfaces destabilized junction-associated ZO-1 and promoted its arrangement into highly dynamic non-junctional cytosolic complexes that averaged ∼2 μm in diameter. Junction-associated ZO-1 rapidly remodeled, and we also observed the direct transformation of cytosolic complexes into junction-like structures. Claudin-family tight junction transmembrane proteins and F-actin also were associated with these ZO-1 containing cytosolic complexes. These data suggest that the cytosolic structures are novel intermediates formed in response to nanotopographic cues that facilitate rapid tight junction remodeling in order to regulate paracellular permeability.

## Introduction

Tight junctions are protein complexes at epithelial apical intercellular contact sites, which form barriers that regulate the paracellular transit of water, ions and molecules (*1-3*). Although their barrier forming properties and morphology suggest a static structure, in fact, tight junction associated proteins are highly dynamic, and can be acutely regulated by external stimuli to alter the extent of paracellular flux (*2, 4*). In particular, the tight junction associated scaffold protein zonula occludens-1 (ZO-1) has been shown to be an important regulator for barrier permeability (*5-9*). ZO-1 is highly mobile, and readily exchanges between tight junctions and the cytosol (*2, 4*). This dynamic process is closely associated with binding to transmembrane tight junction proteins (e.g. claudins, JAM-A) and cytoskeletal proteins, primarily actin (*5, 6, 8*). Detailed understanding of the molecular anatomy of the tight junctions as well as determining how dynamic interactions between tight junction proteins dictate barrier function continue to be elucidated (*2*).

Synthetic materials fabricated with specific geometries and surface topographic features at the micro and nanoscale have the capacity to influence epithelial cell behavior (*10, 11*). We previously found that polymeric films with defined nanostructure, when placed in contact with the apical aspect of an epithelial monolayer, led to enhanced transepithelial permeability of macromolecules ranging in size from ∼60 kDa to ∼150 kDa, including bovine serum albumin and IgG (*12-14*). Immunolabeling of ZO-1 showed characteristic changes in tight junction morphology that were associated with increased permeability, further indicating nanotopography-induced regulation of tight junctions (*12*). Tight junction remodeling occurs relatively quickly (within 1 hour), is reversible, energy-requiring, and depends on MLCK (myosin light chain kinase) signaling (*12, 13*). However, detailing the precise timing and changes to tight junctions that occur in response to nanostructure contact remain to be determined and require specialized methodology. Specifically, commonly used methods to detect epithelial paracellular flux, including transepithelial electrical resistance (TEER) and transepithelial diffusion through porous Transwell inserts (*15, 16*), measure bulk epithelial layer behavior and are unable to obtain submicrometer scale precision (*2*). Moreover, systems incorporating fluorescently tagged proteins-of-interest for live cell imaging are frequently subject to overexpression, which can alter physiological behavior (*1, 15, 17*).

To advance our ability to analyze the effects of nanotopography on epithelial cell permeability, we developed a novel method to achieve high-resolution visualization of paracellular flux across live cell monolayers in real time, using total internal reflection fluorescence (TIRF) microscopy, which specifically images the basal side of epithelia within ∼100 nm from the glass substrate (*18*). We also tagged endogenous ZO-1 protein under the control of its endogenous promoter with mCherry using CRISPR-Cas9 based gene editing, enabling live cell tracking of morphological changes to ZO-1. Through these advanced imaging methods and fluorescence recovery after photobleaching (FRAP) assays, we identified novel nanostructure-induced dynamics of junction-associated ZO-1. This remodeling process was mediated through the formation of cytosolic structures containing ZO-1, claudin-family transmembrane tight junction proteins and F-actin that dynamically interacted with tight junctions. These ZO-1 positive cytosolic complexes are consistent with recently reported structures induced by ZO-1 phase separation that have been implicated as mediators for tight junction formation and mechanosensing (*19, 20*). Our ability to use nanotopography to stimulate active tight junction remodeling by this novel biophysical process provides a basis to understand mechanisms that regulate tight junction assembly and epithelial barrier function in response to the extracellular microenvironment.

## Results

### Nanotopographic cues induce paracellular flux of macromolecules across epithelial monolayers

Human epithelial colorectal adenocarcinoma (Caco-2) cells were cultured on glass-bottomed chambers under conditions that enabled them to form a polarized monolayer (**Fig. 1A**, fig. S1A). The cells were then placed in contact with polypropylene films with or without a defined nanotopographic structure (**Fig. 1B**, fig. S1B). FITC-labeled IgG (FITC-IgG) was added to the apical side of the monolayer as a tracer for barrier permeability (**Fig. 1A**, fig. S1A). We then tracked FITC-IgG that penetrated through epithelial monolayers to the basal side with submicron resolution, using TIRF microscopy that selectively illuminated fluorophores within ∼100 nm zone above the basal substrate (**Fig. 1A**). Acquired TIRF images were quantified as the mean FITC fluorescence intensity at cell-cell borders marked by bulk plasma membrane using Cell Mask Deep Red (fig. S1C). Strikingly, FITC-IgG accumulated in basolateral gaps below cell-cell borders of Caco-2 monolayer after apical contact with nanostructured (NS) films for 1 hour at 37°C (**Fig. 1, C and D**), while FITC-IgG showed minimal paracellular permeability in non-treated (NT) cells (fig. S1 D and E). NS film treated cells also showed significantly higher paracellular accumulation of FITC-IgG than cells in contact with control flat (FT) polypropylene films (**Fig. 1C-E**). These data are consistent with our previous studies, which demonstrate increased paracellular permeability to macromolecules when epithelial cells are in contact with nanotopographic structures (*12-14*), further indicating the regulation of tight junctions through nanostructure contact. We then immunostained ZO-1 in differentially treated cells (fig. S1F) given its important role in tight junction regulation (*5-9*). We found NS-specific effects on ZO-1 morphology, including induction of a zigzag appearance in the xy focal plane (**Fig. 1, F and G**) and xz/yz focal plane (fig. S1, G and H), and interestingly, a novel class of cytosolic complexes (**Fig. 1, F and H**).

**Fig. 1.**
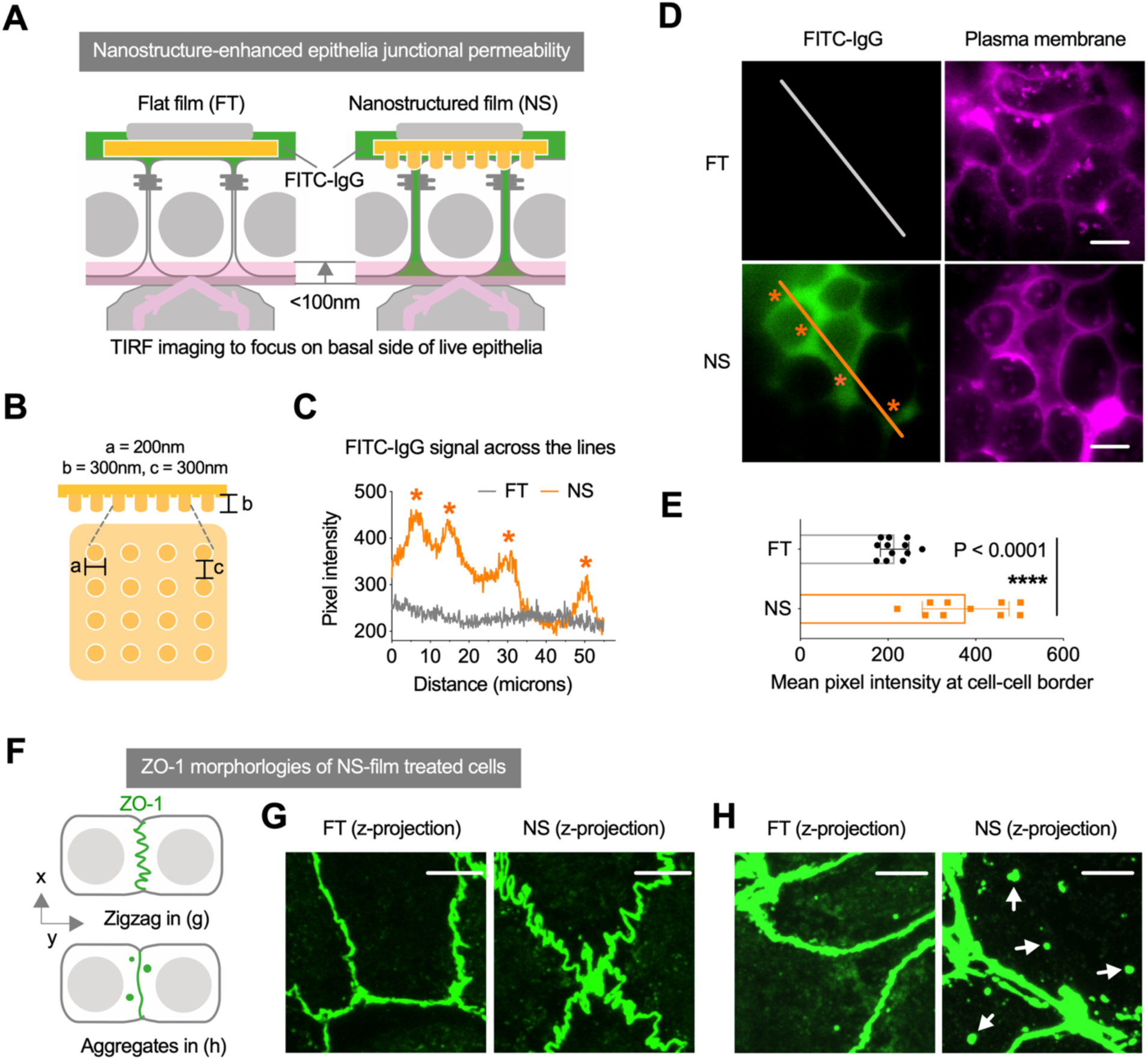
Enhanced permeability of epithelial monolayer barriers through paracellular junctions induced by contact with nanotopographic surfaces. (**A**) Schematic of live epithelial cell basal imaging using total internal reflection fluorescence (TIRF) microscopy to track penetration of fluorescently labeled IgG. (**B**) Schematic of topographic profile of nanoimprinted polypropylene film generated from molds using electron-beam lithography. (**C**) TIRF images of the basal aspect of live Caco-2 monolayers treated with flat (FT) or nanostructured (NS) films together with FITC-labeled IgG (green) applied to the apical side for 1 hour at 37°C (from n = 3 independent experiments). The pattern of FITC-IgG accumulation at the basal side of cells treated with NS films correlated well with basolateral gaps as delineated by cell membrane staining (pink), indicating paracellular permeability. Scale bar: 10 μm. (**D**) Transverse profile of FITC-IgG as indicated by the lines in (C), showing the accumulation of FITC-IgG at basolateral gaps due to NS film treatment. (**E**) Mean fluorescence intensity of FITC-IgG at below cell-cell contact sites in (C). Data are mean ± s.d. (n = 10 images), and the *P* value was determined by two-tailed unpaired *t* test. (**F**) Schematic of ZO-1 morphological changes due to direct contact with NS films. (**G**,**H**) Confocal microscope images of immunostained ZO-1 in Caco-2 cells treated with NS or FT film for 1 hour at 37°C, showing zigzag and aggregate morphologies (white arrows) induced by nanostructures. Images are projections of maximum pixel intensity of z-stack images acquired with 6 μm depth and 0.3 μm intervals. Scale bar: 5 μm.

### Live cell imaging reveals cytosolic tight junction protein complexes induced by contact with nanostructured films

Considering NS-induced morphological changes in ZO-1 (**Fig. 1, F to H**) and its important role in tight junction regulation (*8, 9*), we engineered cells using CRISPR-Cas9 based genome editing to express ZO-1 tagged at the N terminus with a fluorescent reporter (mCherry) for live cell imaging. Using this approach, the ZO-1 was precisely tagged and endogenously expressed, thus maintaining its physiological activity (*21*) (**Fig. 2A**). Briefly, the guide RNA (gRNA) was designed to target exon2 of *TJP1* gene for site specific insertion/deletion (indel) (fig. S2A). Thereafter, the *mCherry* gene with two 1kb arms homologous to the indel site was integrated into the genome through homology directed repair (HDR) (**Fig. 2A**). Transduced Caco-2 cells were selected for mCherry expressing cells through fluorescence-activated cell sorting (FACS) (fig. S2B), followed by single clone isolation and genomic PCR confirmation (fig. S2C). The 19 clones we isolated were all heterozygous with only one allele modified (fig. S2C). Isolated clones cultured on Transwells were used for phenotypic confirmation. Based on TEER analysis, clone15 had barrier function comparable to wildtype cells and thus was used for detailed imaging analysis (fig. S2D). Noticeably, live imaging and z-stack scanning of engineered cells showed the existence of cytosolic mCherry-tagged ZO-1 complexes near tight junctions, mostly adjacent to the apical surface (fig. S2F).

**Fig. 2.**
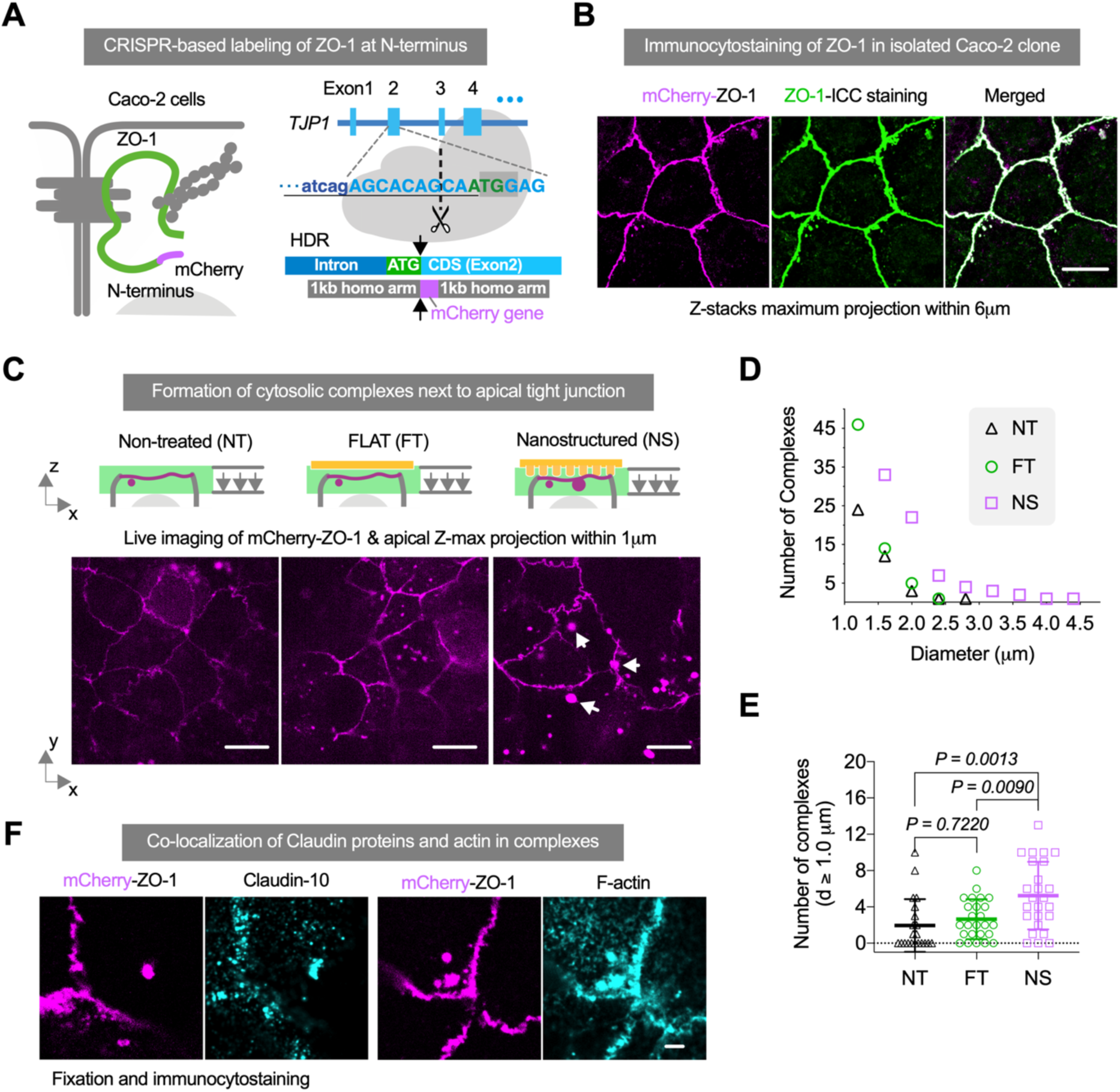
Cytosolic complexes of ZO-1 induced by nanostructures, which also contain claudin-family transmembrane proteins. (**A**) Schematic of CRISPR-based tagging of mCherry reporter to the ZO-1 N-terminus under control of the endogenous promoter. The *mCherry* gene was inserted into exon2 of *TJP1* gene through homology-directed repair (HDR). (**B**) Co-localization of mCherry and immunostained (ICC) ZO-1 in a selected Caco-2 clone validated the fidelity of tagged-ZO-1 for live cell tracking. Images are projections of maximum pixel intensity of apical z-stack images acquired with 1 μm depth and 0.3 μm intervals. Scale bar: 10 μm. (**C**) Images of apical mCherry-ZO-1 in live Caco-2 monolayer cultures in contact with nanostructured (NS) or flat (FT) polypropylene films for 0.5-1.5 hours at 37°C, and non-treated (NT) control. Cytosolic complexes larger than 1.0 μm diameter (white arrows) appeared at the apical side of cells in contact with a NS films. Images are projections of maximum pixel intensity of apical z-stack images acquired with 1 μm depth and 0.3 μm intervals. Scale bar: 10 μm. (**D**) Size distribution of apical cytosolic complexes (d ≥ 1.0 μm) in images represented in (C). (**E**) Number of (d ≥ 1.0 μm) complexes of each image in (C). Data are mean ± s.d. (n = 25 images for each treatment, from 5 independent experiments), and *P* values were determined by one-way ANOVA. (**F**) Immunostained images of mCherry-ZO-1 engineered Caco-2 cells after treatment with NS film at 37°C for 1 hour show co-localization of Claudin-10 and F-actin in cytosolic mCherry-ZO-1 complexes. Scale bar: 2 μm.

Live cells expressing mCherry-ZO-1 at physiological levels were highly susceptible to photobleaching. To minimize this effect, a spinning disk confocal microscope with high imaging speed was used for live cell imaging, where mCherry-ZO-1 fluorescence was collected at an apical 1 μm z-depth with 0.3 μm intervals and z-projected at maximum pixel intensity for each time lapse image (**Fig. 2C**). Physical drift and vibration effects were minimized during live cell imaging using the Nikon Perfect Focus System (PFS). Interestingly, we observed the existence of relatively large cytosolic ZO-1 containing structures (∼2 μm) at the apical side next to tight junctions when cells were in contact with NS films, whereas untreated cells and cells treated with FT films showed fewer and smaller cytosolic ZO-1 complexes (**Fig. 2C**). We quantified (*22*) cytosolic structures in 40 μm x 40 μm apical images from 5 independent experiments (fig. S3A), and found a significantly higher number of large cytosolic ZO-1 complexes (d ≥ 1 μm) in NS film-treated cells compared to untreated or FT film-treated cells (**Fig. 2, D and E**). To investigate their composition, we immunostained differentially treated cells for several markers, including ZO-1, claudins, F-actin and the endocytosis marker Rab5. mCherry positive apical cytosolic structures were confirmed to be ZO-1 positive, and also co-localized with several claudins (Claudin-2, −4, −10) and F-actin (**Fig. 2F**, fig. S3, B and C). Notably, the structures were negative for Rab5 staining (fig. S3C), and were non-acidic based on a lack of LysoTracker staining (fig. S3D), indicating that they did not originate from vesicular mediated trafficking of endocytosed tight junction proteins.

### Enhanced dynamics of tight junction remodeling stimulated by nanotopography

We investigated the dynamics of mCherry-ZO-1 remodeling, using time-lapse imaging of Caco-2 cells stimulated by nanotopography. We observed two patterns of cytosolic structure initiation: i) the direct transition of junction-associated ZO-1 into cytosolic structures through aggregation and departure (**Fig. 3A-i**); ii) the clustering of diffused signals into puncta within a network next to tight junctions (**Fig. 3A-ii**, see movie S1). In addition, the cytosolic structures became enlarged through fusion of newly formed small structures (**Fig. 3A-iii**). Interestingly, cytosolic structures were highly dynamic in that they were observed to circulate apically while collecting more ZO-1 followed by movement toward the basal aspect of the cytoplasm (**Fig. 3A-iv**, see movie S2 and movie S3). On the other hand, we also observed the rapid reorganization of ZO-1 into junction-like structures that often originated from non-apical locations (**Fig. 3A-i,v**). Occasionally, we saw direct conversion of spherical structures into junction-like structures (**Fig. 3A-vi**). With TIRF imaging, we also observed ZO-1 positive junction-like structures at the basal side of Caco-2 monolayer stimulated by apical contact with nanotopography (fig. S4A). The appearance of basal ZO-1 positive strands suggests that apical contact with NS films may induce cells to reorient their apical-basolateral axis.

**Fig. 3.**
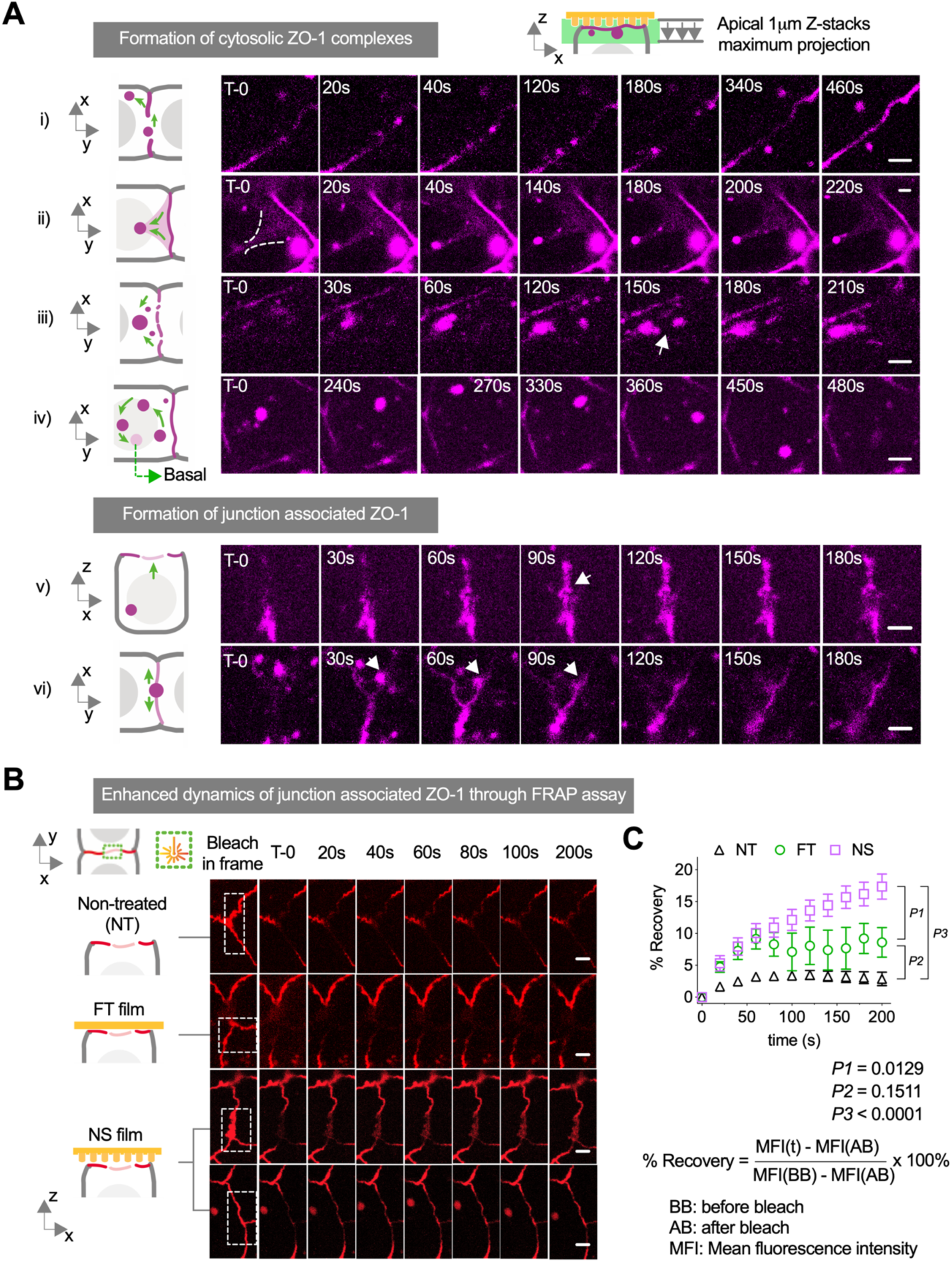
Enhanced dynamics of junction-associated ZO-1 in engineered Caco-2 cells in contact with nanostructured film. (**A**) Time-lapse images of mCherry-tagged ZO-1 at the apical side of the cells stimulated by nanotopographic cues for 0.5-1.5 hour at 37°C. Means of cytosolic complexes formation included: i) junction-associated ZO-1 proteins aggregate and dissociate from the original structure into the cytosol; ii) cytosolic ZO-1 proteins cluster into puncta within a network (highlighted by white dashed line) next to junctions; iii) cytosolic structures merge with newly restructured ZO-1; iv) apical cytosolic complex circulates while collecting smaller ZO-1 structures which then moves basolaterally. Junction-like ZO-1 structure formed through v) rapid reorganization likely originated from basolateral side; and vi) direct transformation of cytosolic ZO-1 complex (white arrow). Images are projections of maximum pixel intensity of apical z-stack images acquired with 1 μm depth and 0.3 μm intervals. Scale bar: 2 μm. (**B**) Time-lapse images of junction-associated mCherry-ZO-1 after photobleaching at a selected region (white dashed line) in response to contact with either a flat (FT) or nanostructured (NS) film for 0.5-1.5 hour or that were non-treated (NT). Scale bar: 2 μm. (**C**) Quantification of fluorescence recovery after photobleaching (FRAP) in selected frames represented in (B). Data are mean ± s.d. (analyzed from 26 images for NT, 11 images for FT, and 29 images for NS, pooled from 2 independent experiments), and *P* values were determined by one-way ANOVA analysis of recovery at 200 s.

In contrast, we did not observe active ZO-1 remodeling in cells that were either untreated or in contact with FT films, where instead there was minimal interaction between tight junctions and apical cytosolic structures, Moreover, there were fewer and smaller ZO-1 positive cytosolic complexes in cells that were not NS-treated (fig. S4, B and C).

We then used FRAP at selected regions to quantify and compare the remodeling rate of tight junctions in response to NS contact (**Fig. 3B**). Importantly, we found that mCherry-ZO-1 fluorescence at tight junctions recovered faster after photobleaching in cells stimulated by NS films, as compared to untreated cells and FT film treated cells (**Fig. 3C**). This further supports that there is enhanced remodeling of junctional ZO-1 being induced by nanotopographic cues.

To examine the dynamics of a transmembrane tight junction protein in live cells, we transfected mCherry-ZO-1 expressing Caco-2 cells with exogenous YFP-claudin-3 using an adenovector expression construct (*23*) (**Fig. 4A**). The virus titer was optimized to minimize YFP-claudin-3 overexpression although this did result in heterogeneous expression by different cells in the monolayer (**Fig. 4B**). YFP-claudin-3 and mCherry-ZO-1 co-localized at apical cytosolic structures in NS film-treated cells (**Fig. 4B**), but not untreated cells or FT film-treated cells (fig. S5). Importantly, claudin-3 was present in ZO-1 containing cytosolic structures at sites of formation and sites of interaction with tight junctions (**Fig. 4, C and D**, see movie S4). The co-localization of the transmembrane tight junction protein claudin-3 with cytosolic scaffold protein ZO-1 indicates that cytosolic complexes involve the regulation of multiple classes of tight junction proteins, including transmembrane proteins.

**Fig. 4.**
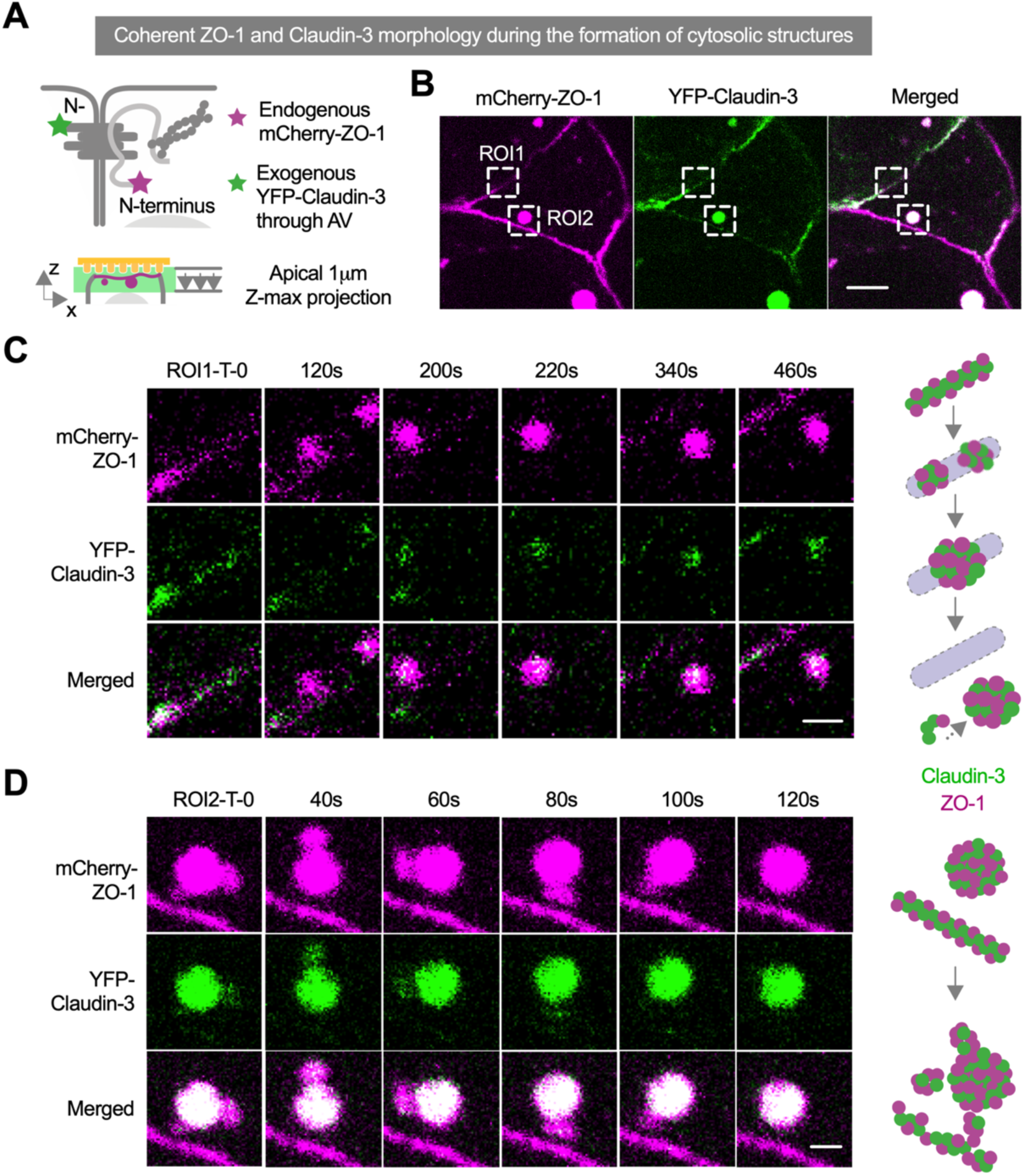
Co-transport of ZO-1 and claudins during the initiation and formation of cytosolic complexes after nanotopographic contact. (**A**) Schematic of exogenous YFP-claudin3 adenovirus transduction in mCherry-ZO-1 engineered Caco-2 cells for live imaging analysis. Images of each time lapse were acquired at apical 1 μm depth with 0.3 μm intervals and projected the maximum pixel intensity into presented images. (**B**) Images of mCherry ZO-1 and YFP-claudin-3 co-expressing Caco-2 cells treated with a nanostructured film at 37°C for 0.5-1.5 hour, showing co-localization of ZO-1 and claudin-3 in apical cytosolic complexes. Scale bar: 10 μm. (**C**) Time-lapse images of ROI1 in (B) show the involvement of both ZO-1 and claudin-3 in the initiation of complex formation and cytosolic migration. (**D**) Time-lapse images of ROI2 in (B) show formation of apical cytosolic complexes containing both ZO-1 and claudin-3 originating from a neighboring tight junction. Scale bar: 2 μm.

### F-actin engages during the formation of cytosolic tight junction complexes

ZO-1 interconnects transmembrane junction proteins (e.g. claudins) and cytoskeletal F-actin in epithelial cells (*5, 6*). We imaged F-actin labeled with SiR-actin in live cells under NS and FT film treatment, to investigate the changes in cytoskeletal morphology in response to different stimuli (**Fig. 5A**). We observed apparent clustering of F-actin in cells upon NS film treatment as opposed to untreated cells or FT film treated cells. These clusters formed in response to NS contact translocated from the apical aspect of the cells to the basal side over 1.5 hour (fig. S6A). More specifically, F-actin interacted with ZO-1 proteins during tight junction remodeling in different ways (see movie S5): i) F-actin replaced sites of ZO-1 protein localization at tight junctions (**Fig. 5B-i**); ii) F-actin intertwined with ZO-1 in cytosolic complexes sitting next to tight junctions (**Fig. 5B-ii**); iii) polarized cytosolic F-actin-ZO-1 structures (**Fig. 5B-iii**) with the F-actin portion facing towards iv) larger solid integrated structure (**Fig. 5B-iv**); and v) F-actin and ZO-1 co-localized in large (>2 μm) hollow structures (**Fig. 5B-v**). In some instances, F-actin:ZO-1 containing structures merged into larger co-localized structures (**Fig. 5C**). And interestingly, there were instances where F-actin dissociated from ZO-1 containing cytosolic structures (**Fig. 5, D and E**). Consistent with this, cytosolic structures containing both ZO-1 and Claudin-3 lacked F-actin (**Fig. 5, F and G**), indicating that interactions of F-actin with cytosolic complexes are transient and F-actin dissociates as the complexes mature.

**Fig. 5.**
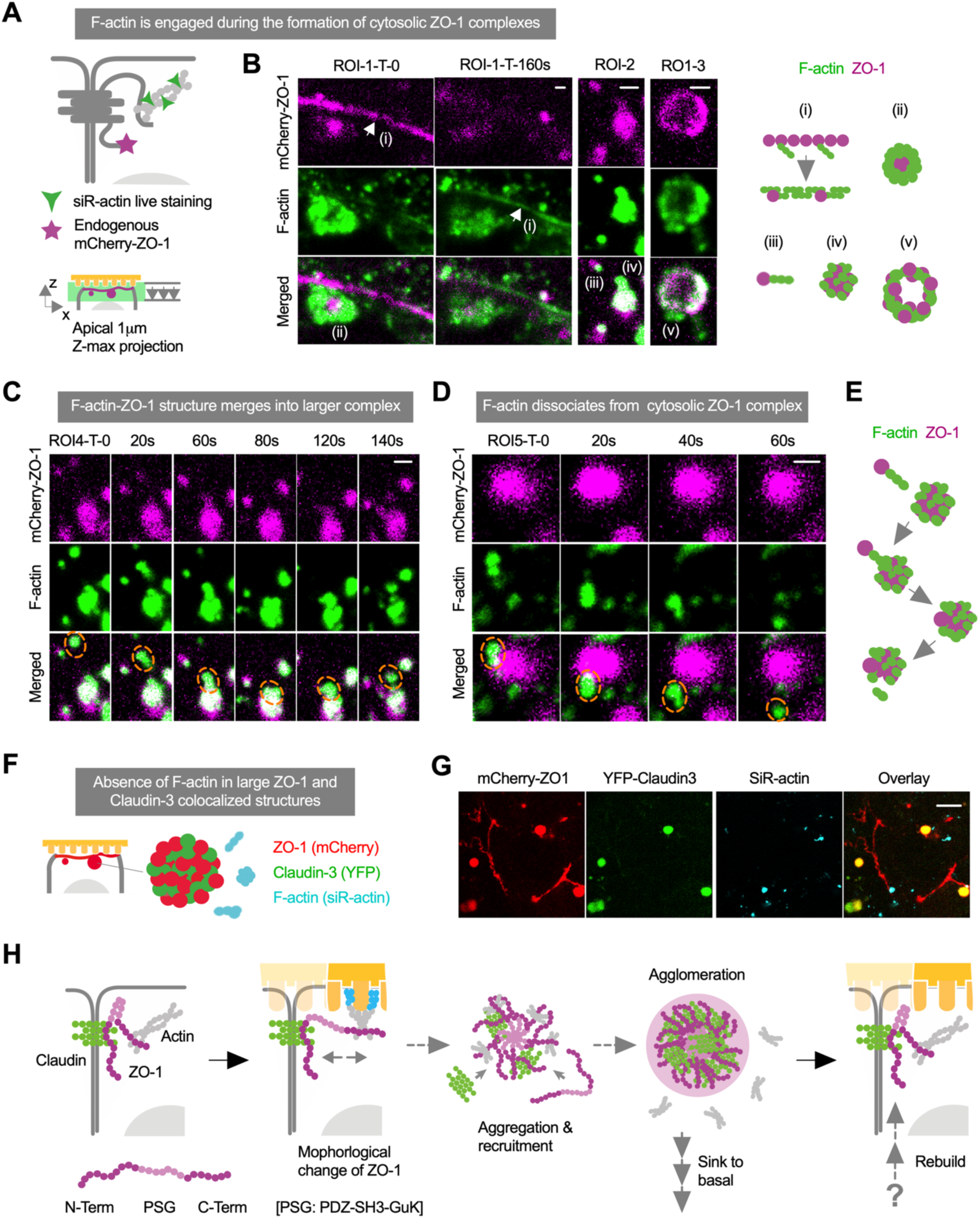
Nanotopographic modulation engages F-actin in the formation of ZO-1 cytosolic complexes. (**A**) Schematic of F-actin staining in mCherry-ZO-1 engineered Caco-2 cells for live imaging analysis. Images of each time lapse are acquired at apical 1 μm depth with 0.3 μm intervals and projected the maximum pixel intensity into presented images. (**B**) Images of cells stimulated by nanotopographic cues for 0.5-1.5 hour at 37°C show different patterns of F-actin/ZO-1 interactions: i) disappearance of ZO-1 concomitant with the appearance of F-actin at tight junctions; ii) F-actin wrapped around ZO-1 in cytosolic complexes next to a tight junction; iii) polarized cytosolic F-actin-ZO-1 complexes with F-actin facing toward iv) a larger solid integrated structure; and v) organization of F-actin and ZO-1 into large (>2 μm diameter) hollow structures. Scale bar: 2 μm. (**C**) Time-lapse images show merging of the F-actin-ZO-1 structure (orange dashed line) into a larger complex containing the two proteins. Scale bar: 2 μm. (**D**) Time-lapse images show dissociation of F-actin from cytosolic ZO-1 complex. Scale bar: 2 μm. (**E**) Summary schematic of F-actin associated dynamics involved in formation of cytosolic ZO-1 complexes. (**F**) Schematic of mCherry-ZO-1, YFP-claudin-3 and F-actin interactions in engineered Caco-2 cells for live imaging analysis. (**G**) Images of live cells that are stimulated by nanotopographic cues for 72 mins at 37°C show co-localization of ZO-1 and claudin-3 but not F-actin in apical cytosolic complexes. Scale bar: 10 μm. (**H**) Proposed mechanism of tight junction remodeling in response to nanotopographic cues involving ZO-1 cytosolic complexes.

Taken together, our data support the hypothesis that NS materials in direct contact with the apical surface of epithelial cells induce F-actin rearrangements. The rearrangements are coordinated with changes in localization and morphologies of its binding partner, ZO-1. Transition of junction-associated ZO-1 into large cytosolic complexes also recruit other tight junction proteins, suggesting a new mechanism of tight junction regulation by biophysical stimuli (**Fig. 5H**).

## Discussion

Tight junctions, comprised of heavily cross-linked complexes of transmembrane and membrane-associated proteins, represent a structural determinant of epithelial cell polarity and regulate solute paracellular permeability (*1, 3*). Tight junctions are highly dynamic, even in unstimulated cells, and are acutely regulated by multiple extracellular biological stimuli including hormones and cytokines (*2, 4*). We previously found *in vitro* and *in vivo* that cell contact with polymeric films with defined nanotopographic features enhances transepithelial permeability to macromolecules in the size range of (60 kDa to 150 kDa) (*12-14*). Here, for the first time, we used state-of-the-art TIRF microscopy to visualize paracellular flux of FITC-labeled IgG with submicron resolution at the basal aspect of live epithelial monolayers stimulated with a nanostructured biophysical cue on the apical surface. The ability to image transepithelial flux in live cell monolayers provided more detailed morphological information than methods measuring bulk solute flow and enabled us to demonstrate that the paracellular route is a major pathway for nanostructure-induced transepithelial permeability.

ZO-1 has been shown to play an essential role in the maintenance and regulation of tight junction permeability (*8, 9, 20*). To track the intracellular distribution of ZO-1 in live cells, we used CRISPR-Cas9 based gene editing to attach a fluorescent mCherry reporter to the N-terminus of endogenous ZO-1. Engineered ZO-1 expression was driven by its endogenous promoter which ensured that it was expressed and regulated at physiologically relevant concentrations avoiding pitfalls related to overexpression (*1, 19*). Live imaging and FRAP analysis of cells triggered by nanostructured films revealed a change in the kinetics of ZO-1 turnover associated with the response of tight junctions to the contact between cells and NS films. Our findings also are consistent with previous studies demonstrating that tight junction associated ZO-1 is highly mobile and exchanges with the cytosolic pool (*2, 4*). Furthermore, the imaging methods used here also enabled us to identify a novel intermediate involved in nanostructure-induced remodeling tight junctions, namely, large (∼2 μm diameter) cytosolic ZO-1-containing complexes that co-localize with claudin-family proteins and, in some cases F-actin. This ZO-1 positive cytosolic structure has the characteristics of a newly described complex that forms as a result of liquid-liquid phase separation of ZO-1 (*19*). Phase separation of ZO-1 and its accompanied junctional proteins was also reported as an important intermediate for tight junction assembly (*19*) and mechanosensing (*20*). Here we extend these observations by showing that a nanostructured surface with the capacity to increase transepithelial permeability also induces the formation of ZO-1 positive cytosolic complexes in conjunction with an increase in tight junction remodeling. In addition, the previous studies demonstrated the formation of liquid-liquid phase separated ZO-1 complexes in cell free systems and cells overexpressing ZO-1 (*19*). Critically, the engineered cells we examined expressed comparable, physiologic levels of mCherry-ZO-1 as untagged ZO-1, thus demonstrating the formation of these complexes can occur in response to a physiologic stimulus and was not driven by protein overexpression.

ZO-1 is a peripheral tight junction protein that interconnects transmembrane tight junction proteins (e.g. claudins, occludin, JAM) and F-actin (*5, 6*). Based on structure-function analysis, ZO-1 phase separation was attributed to the unfolding of the PSG (PDZ-SH3-GuK) domain, which is directly regulated by binding to the F-actin cytoskeleton (*7, 9, 19*). Moreover, it has been previously shown that ZO-1 positive cytosolic complexes are associated with the actin cytoskeleton that facilitated their delivery to nascent tight junctions in response to changes in tension that occur during gastrulation (*20*). Consistent with this model, we also observed that F-actin transiently interacted with cytosolic structures in response to nanotopography. A role for tension in the formation of ZO-1 cytosolic complexes is also suggested by the observation that integrin stimulation is required for nanotopography-induced epithelia permeability (*12, 13*). We also found that ezrin, a cytosolic scaffold protein that links the actin cytoskeleton to cellular membrane receptors (*24*), was upregulated in NS-treated cells (fig. S6B). Thus, a signal transduction pathway from membrane receptors to ezrin, F-actin, ZO-1, and then to other junctional proteins, may transduce physical signals from cell apical contacts to modulate tight junction function. This agrees with the recent work demonstrating that ZO-1 phase separation mediates mechano-sensing of tight junctions (*20*).

In summary, we have developed a series of advanced imaging methods that enabled the visualization of transepithelial penetration at submicron resolution and the analysis of tight junction protein dynamics in live cells. Using these approaches, we found that there were hallmark changes in ZO-1 trafficking that were associated with increased paracellular permeability of macromolecules induced by nanotopographic cues. Of particular interest, our data demonstrate that unique ZO-1 containing cytosolic complexes can be induced to interact with tight junctions to regulate their permeability. Our data also underscore that nanotopography is a unique stimulus for fast and reversible modulation without permanent loss of epithelial monolayer integrity. In addition, the imaging techniques shown here have general applicability to understanding the mechanistic basis for tight junction permeability and how barrier function can be modulated by other stimuli in real time at the level of individual intercellular junctions.

## Materials and Methods

### Cell lines and culture

Caco-2 human colon epithelial cell line was purchased from ATCC (#HTB-37). Unless specified, cells were cultured in DMEM (Sigma, #D5796) supplemented with 20% FBS (Gemini Bio #100-106), 100 mM sodium pyruvate (Sigma #113-24-6), and penicillin-streptomycin (Sigma #516106) incubated in 5% CO_2_ at 37°C. The cells were subcultured at 90% confluency by trypsinization with 0.25% trypsin-EDTA (SM-2003). For transepithelial electrical resistance (TEER) measurements, cells were seeded in a 6.4 mm Transwell inserts (Corning #353495) with 300 μL medium containing 84,000 cells/cm^2^ seeded in the upper chamber and 500 μL medium in the bottom chamber. The medium was replenished every other day, and TEER was measured using a voltohmmeter (World Precision Instruments) from day 6-16 where the measured resistance in Ohms was multiplied by the area of the Transwell filter (0.3 cm^2^). For live cell imaging, 200,000 cells were seeded on a 35 mm glass bottom dish with 14 mm micro-well #0 cover glass (Cellvis #D35-14-0-N), which was precoated with 0.3 mg/mL Matrigel (Corning #354234) at 37°C for 30 minutes. Cell culture medium was replenished every other day. After 8-10 days, the monolayer was exchanged with cell imaging medium containing FluoroBrite DMEM Media (Thermo #A1896701) and 20% FBS for live cell imaging analysis.

### CRISPR/Cas9 knock-in in Caco-2 cells

To generate an N-terminal mCherry knock-in in the initial exon of ZO-1, guide RNA (gRNA) for CRISPR/Cas9 mediated site-specific gene editing was designed using Benchling CRISPR tool (www.benchling.com). gRNA targeting *TJP1* gene exon number 2 - 5’CCTTTATCAGAGCACAGCAA3’ was synthesized and complexed with trans-activating crRNA by IDT. 500,000 Caco-2 cells were suspended in 100 μL buffer (Lonza #VCA-1002) supplemented with 264 pmol gRNA duplex, 10 μg Cas9 expression plasmid (GE Healthcare #U-005100-120), and 10 μg repair plasmid (Genscript, Note S1), followed by electroporation using program B024 of Nucleofector™ 2b Device (Lonza #AAB-1001). Cells were then plated in one well within a 24-well plate supplemented with 1 mL medium. To validate CRISPR/Cas9 and gRNA mediated site-specific insertion/deletion (indel), cells from the transfection without repair plasmid were collected after 3 days for genome extraction (Bioline #BIO-52066) and PCR (NEB #M0491S) at the region of indel (forward primer: 5’TGTTTGTGACGTTAAAGCAGCC3’, reverse primer: 5’CACAAACTTACCCTGTGAAGCG3’). Genome indel was assessed by T7 Endonuclease I assay (NEB #M0302) followed by SDS-PAGE gel electrophoresis (Genscript #M42012L). Seven days after the transfection of the complex including the repair plasmid, cells were collected and sorted by FACS (Sony SH800) for enrichment of mCherry positive cells. Enriched cells were plated and subcultured in 10 cm dishes at 3000 cells per dish to produce single cell clones. Nineteen clones were isolated after 2 weeks, further expanded, and confirmed for mCherry knock-in through genome extraction, PCR (using the same primer pair as above) and analyzed by agarose gel electrophoresis.

### Nanostructured film fabrication and cell treatment

Nanostructured films were made, as previously described (*12-14*), by nanoimprint lithography in which polypropylene was heated above its glass transition temperature and pressed into a silicon mold. Molds were fabricated using electron beam lithography followed by anisotropic reactive ion etching to generate precise submicron structures. Nanostructured films were characterized using a Phenom scanning electron microscope (SEM). Nanostructured films or flat (unstructured) films (outside of the imprinted region) were biopsy punched into 6 mm diameter circles, the backside of which was glued to a polyethylene terephthalate film with a 0.3-inch of pipette tip attached. This device was placed in direct contact with a cell monolayer on the apical side and precision weighted with two metal rings of ∼0.2 g and used for cell studies.

### TIRF microscopy for paracellular flux analysis

Caco-2 cell monolayers, cultured on glass-bottom tissue culture dishes, were stained for plasma membranes (CellMask Deep Red Stain, 5 μg/mL, Thermo #C10046) and nuclei (Hoechst 33342, 5 μg/mL, Thermo #H3570) at 37°C for 10 mins in DMEM medium without FBS, followed by two rinses using cell imaging medium. An NS (nanostructured) or FT (flat) film device was placed in contact with cells on apical side along with FITC-IgG (Sigma #F9636) supplemented to the cell imaging medium at 10 μg/mL. After 1 h incubation at 37°C, live cells were imaged using an OMX-SR microscope (GE Health Care) in the ring-TIRF mode equipped with PCO Edge 5.5 cMOS cameras, 4-line laser launch 405/488/568/60nm (Toptica), and live cell chamber at 37° with 5% CO_2_. Images were acquired using a Plan ApoN 60x/1.42 (Olympus) oil immersion objective with laser liquid 1.518 (Cargile), and filter sets for DAPI (435/31), GFP (528/48), mCherry (609/37) and Cy5 (683/40). Registration alignment was determined using an image registration target slide (GE Health Care, pat #52-852833-000) and processed with SoftWoRx 7.0.0. software (GE Health Care). To quantify paracellular permeability of FITC-IgG, cell-cell border area in each image was outlined by a mask, generated using signals from plasma membrane stain, using Fiji software. Mean pixel intensity of FITC signal was analyzed within mask-identified area.

### Confocal microscopy of live Caco-2 cells

Caco-2 cells were fluorescently labeled for different proteins and cell machineries as specified before live cell film device treatment and imaging. For fluorescence tagging of claudin-3, adenovirus carrying YFP-claudin-3 gene were prepared as described (*23*), and added to Caco-2 cells at ∼100 MOI (multiplicity of infection) 2 days prior to imaging. Adenovirus containing medium was removed by fresh medium exchange one day after infection. For F-actin staining, SiR-actin (Cytoskeleton, #CY-SC001) was added into the medium at 0.2 μM the day before imaging, and the medium was replenished with the imaging medium right before imaging. For lysosome staining, LysoTracker Blue (Invitrogen #L7525) was added to the cell culture medium at 60 nM and incubated for 1 h at 37°C, followed by the imaging medium replenishment prior to imaging. Live cell images were acquired within 90 minutes after the treatment of film device, on a spinning disk confocal microscope (Nikon Ti inverted microscope with Andor Borealis CSU-W1 spinning disk and Andor Zyla 4.2 sCMOS camera) equipped with Plan Apo VC 100x/1.4 (Olympus) oil immersion objective, 4-line laser launch 405/488/561/640nm (Andor), and live cell chamber at 37° with 5% CO_2_. The microscope was controlled using micro-manager, and the Nikon Perfect Focus System was used to adjust for axial focus fluctuations.

### Fluorescence recovery after photobleaching (FRAP)

FRAP experiments in cells were carried out using the Nikon spinning disk confocal microscope described above with the following settings. Each regions of interest (ROI), less than 10 μm x 10 μm, was bleached using a 473 nm diode with 30 mW at the back focal plane of the objective with 10 repeats. Several junctions within the field of view were bleached but only one junction per cell. Pre-bleach and post-bleach images were acquired with a 561 nm laser with 30 mW and 300 ms exposure. Fluorescence recovery of mCherry was monitored for 200 s with a time resolution of 20 s. Mean fluorescence intensity (MFI) of the bleached area was quantified by Fiji software, and cell movements during the recovery were corrected by manual adjustment. Image groups were blinded for the analyzer during the quantification. Relative FRAP efficiency was calculated using the following formula (t: time point, BB: before bleach, AB: after bleach), and compared among different treatment groups: NT (non-treated), FT (flat), and NS (nanostructured).

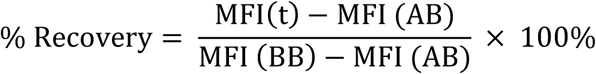

### Quantification of cytosolic complexes

A 40 μm x 40 μm frame that covers cell apical side with tight junctions in each image was selected and the cytosolic ZO-1 complexes were quantified using ImageJ with following steps (*22*). The image was first applied a median-blur filter to reduce noise, followed by segmentation of cytosolic structures using the interactive h-maxima watershed tool, where parameters including seed dynamics, intensity threshold, and peak flooding, were optimized and kept constant for images across treatments within the same independent experiment. The binary image generated from the watershed results included the majority of cytosolic structures and a portion of tight junctions as well. Thereafter, the particle analysis function was used to isolate spherical cytosolic structures and report size distribution. Image groups were blinded for the analyzer during the quantification.

### Immunostaining

Caco-2 cells on tissue culture dishes were rinsed with PBS (with Ca^2+^ and Mg^2+^) three times, and fixed at room temperature in 2% paraformaldehyde/PBS (Electron Microscopy Sciences #RT-15710) for 15 minutes. Cells were then washed once with PBS and residual paraformaldehyde was quenched using 1M glycine (Sigma #50046) in PBS for 10 minutes. Each of the following wash steps were carried out at room temperature for 5 minutes. After three washes using PBS, cells were further fixed/permeabilized using methanol/acetone solution (v/v:50/50) for exactly 2 minutes at room temperature. Cells were then sequentially washed three times with PBS, once with 0.5% Triton™X-100 in PBS, and twice with 2% goat serum (Sigma #G9023) and 0.5% Triton™X-100 in PBS. Primary antibodies including anti-ZO-1 (Thermo #339188), anti-Claudin-2 (#ab53032), anti-Claudin-4 (Abcam #53156), anti-Claudin-10 (Thermo #38-8400), anti-Rab5 (Abcam #ab18211) and anti-Ezrin (Abcam #ab4069) were used for specific immunostaining, and cells were stained for 1 hour at room temperature in PBS with 2% goat serum and antibody supplemented, followed by three washes using PBS with 2% goat serum. Cells were then stained with secondary antibodies including goat anti-mouse-Alexa Fluor 488 (Thermo #A11029) and goat anti-rabbit-Alexa Fluor 633 (Thermo #A21070) in PBS with 2% goat serum at room temperature for 1 hour. For F-actin staining after fixation, Alexa Fluor 488 Phalloidin was added into the secondary antibody staining mixture. Cells were washed three times with PBS followed by imaging using the spinning disk confocal microscope described above.

### Statistical analysis

All data are expressed as the mean ± the standard deviation (SD). The value of n and what n represents (e.g., number of images, experimental replicates, or independent experiments) is stated in figure legends and results. Statistical analysis was performed in Prism 8 (GraphPad Inc.), and the statistical test used is indicated in the relevant figure legend.

## General

We thank K. Herrington for advice on cell imaging.

## Funding

this research was supported by the National Institute of Health grant 1R01EB018842 to T.A.D. and M.K. X.H. was supported by a Li foundation fellowship. X.S. was supported by the NIH Pathway to Independence Award 1K99GM126136. B.H. is a Chan Zuckerberg Biohub Investigator.

## Author contributions

X.H., X.S., M.K., T.A.D designed the research. X.H. carried out the experiments. X.H. and M.E.H analyzed the data. X.H., X.S., A.C., T.M., M.K. and T.A.D interpreted the results. X.H. drafted the manuscript. X.H., M.K., X.S., C.N., B.H., T.M. and T.A.D. edited the manuscript.

## Competing interests

the authors declare no competing interest.

## Data and materials availability

All data needed to evaluate the conclusions in the paper are present in the paper and/or the Supplementary Materials. Additional data related to this paper may be requested from the authors.

## Supplementary Materials

**Fig. S1.**
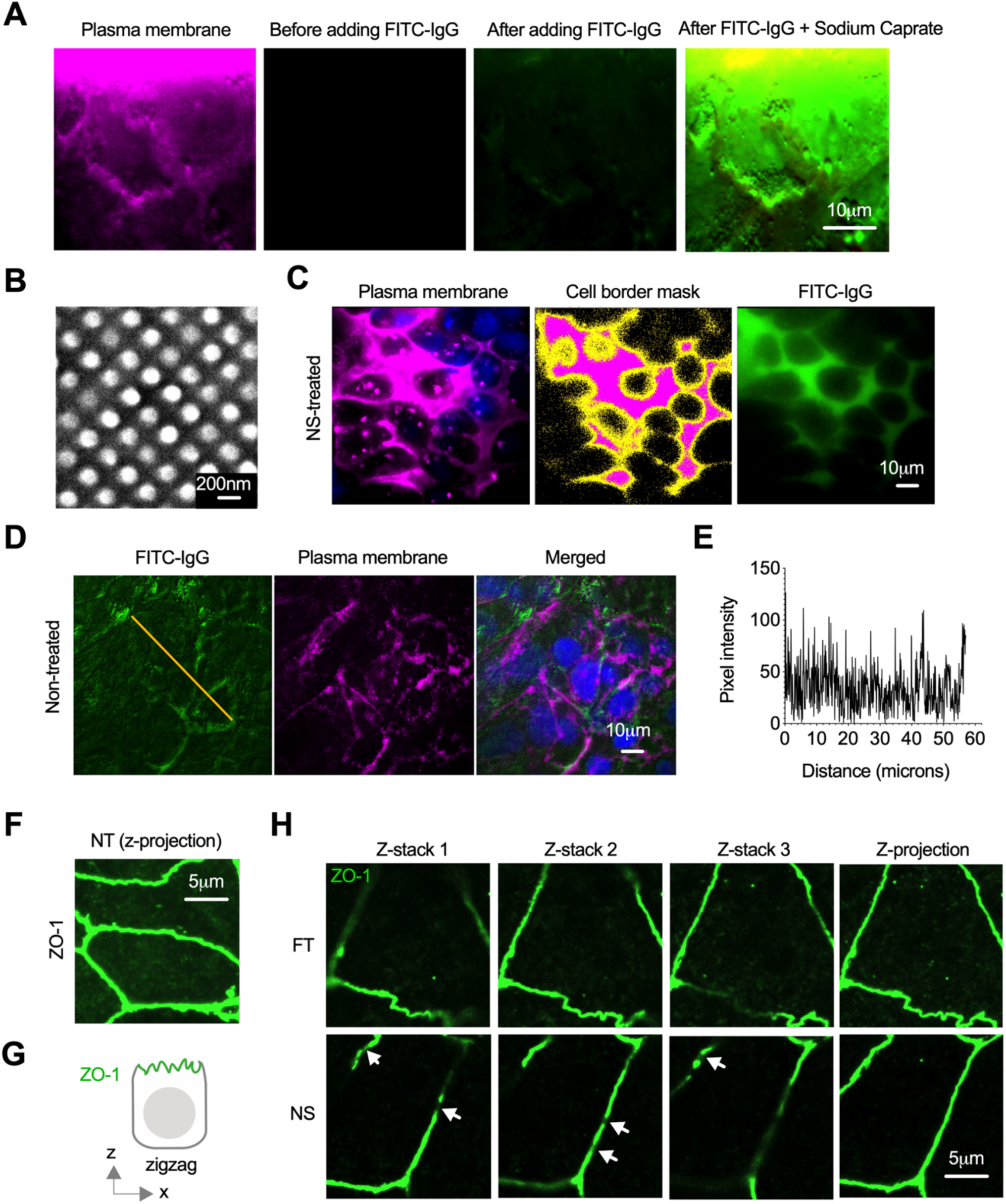
Enhanced leakiness of biomolecules through paracellular junctions with nanotopographic film treatment. (**A**) TIRF images of basal side of live Caco-2 monolayer treated with FITC-labeled IgG and sodium caprate, showing polarized epithelial monolayer with paracellular barriers blocking the penetration of macromolecules from apical side to basal side. Adding sodium caproate at 1 mM irreversibly opened tight junctions and led to immediate penetration of FITC-IgG through the monolayer to be detected by TIRF imaging at the basal side. (**B**) Scanning electron microscopy (SEM) image of nanostructured polypropylene film. (**C**) Images of cell-cell border area mask generation from plasma membrane staining, which is used for quantification and normalization of FITC-IgG signal accumulation. (**D**) TIRF images of basal membrane of live epithelial cell Caco-2 monolayer treated with FITC-labeled IgG for 1 hour at 37°C (from n = 3 independent experiments). (**E**) Transverse profile of FITC-IgG as indicated by the lines in (D). Signal intensity and distribution in FITC-IgG channel is not comparable to the conditions where the flat/nanostructured film are in direct contact with cells (Fig. 1c) due to the reflection effect of the film during TIRF imaging. (**F**) Image of immunostained ZO-1 in non-treated Caco-2 cells, as compared to Fig. 1g&h. (**G**) Schematic of ZO-1 zigzag morphology at xz/yz focal plane of Caco-2 cells in direct contact with nanostructured film. (**H**) Images of immunostained ZO-1 in Caco-2 cells treated with NS or FT film for 1 hour at 37°C show zigzag morphology at xz/yz focal plane. Z-projection images are projections of maximum pixel intensity of z-stack images acquired with 6 μm depth and 0.3 μm intervals.

**Fig. S2.**
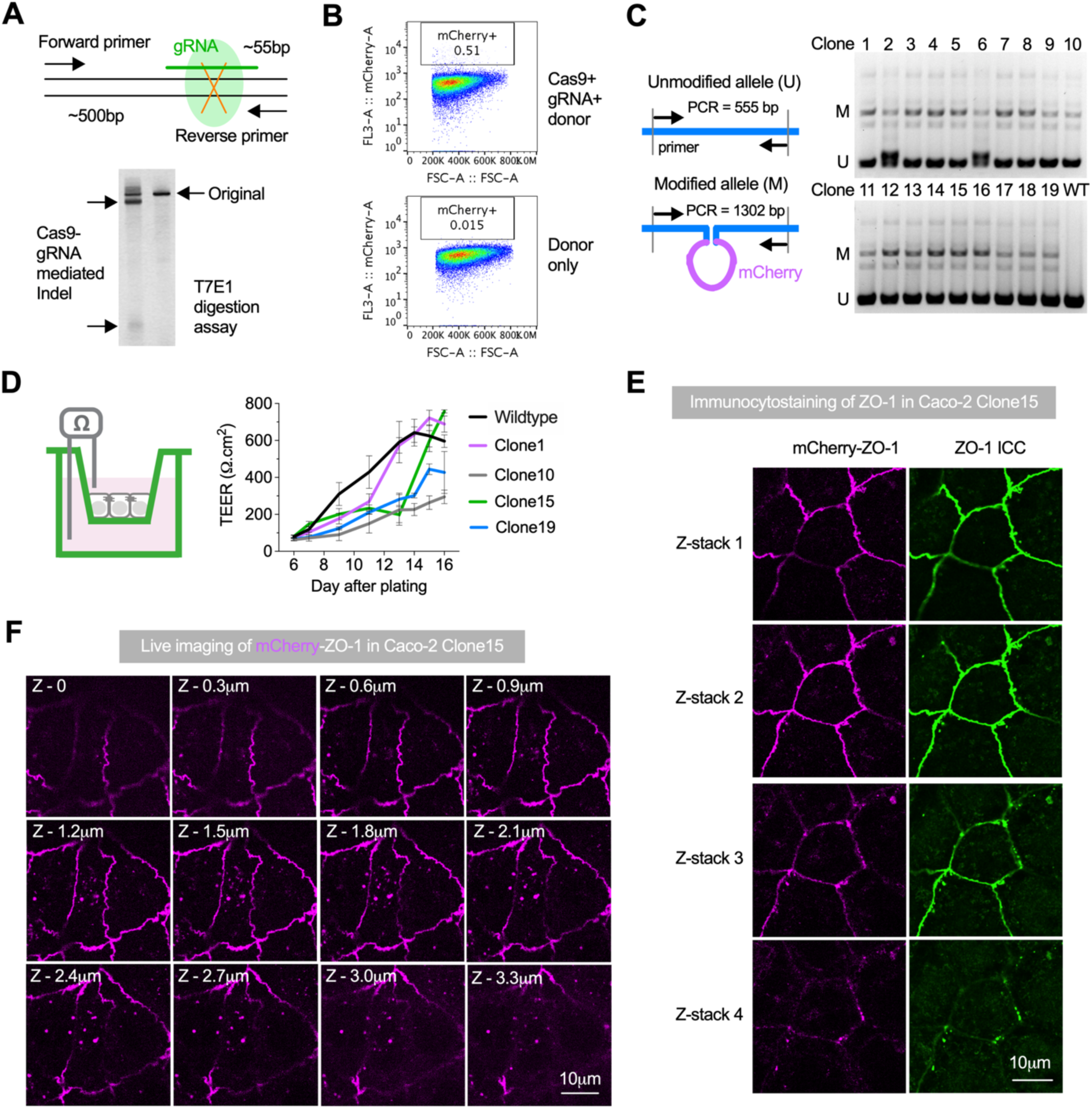
Tagging of endogenous ZO-1 with mCherry reporter in Caco-2 cells through CRISPR-based genome editing and its phenotypic confirmations. (**A**) T7E1 digestion assay and agarose gel electrophoresis to detect Cas9-gRNA mediated insertion/deletion (indel) of targeted *TJP1* gene. (**B**) FACS sorting of CRISPR-transduced Caco-2 cells to enrich mCherry positive population (in frame). Top panel: forward scattering and mCherry signal plot of cells co-transfected with Cas9, gRNA and donor template DNA after 14 days; bottom panel: forward scattering and mCherry signal plot of un-transfected control cells. (**C**) Genomic PCR and agarose gel electrophoresis of purified clones to confirm the insertion of mCherry gene at intended loci. All of the isolated clones have heterozygous modification of ZO-1. (**D**) Validation of barrier formation potency of engineered Caco-2 clones through transepithelial electrical resistance (TEER) measurements of cells cultured on transwell inserts. Clone15 with optimal accumulation of TEER values was selected for imaging analysis. Data are mean ± s.d. (n = 3 biological replicates). (**E**) Co-localization of mCherry and immunocytostaining (ICC) of ZO-1 in Caco-2 Clone15 at different Z-stacks. (**F**) Z-stack scanning of live Caco-2 Clone15 by confocal microscopy reveals cytosolic structures of ZO-1 at basal-lateral locations underneath tight junctions.

**Fig. S3.**
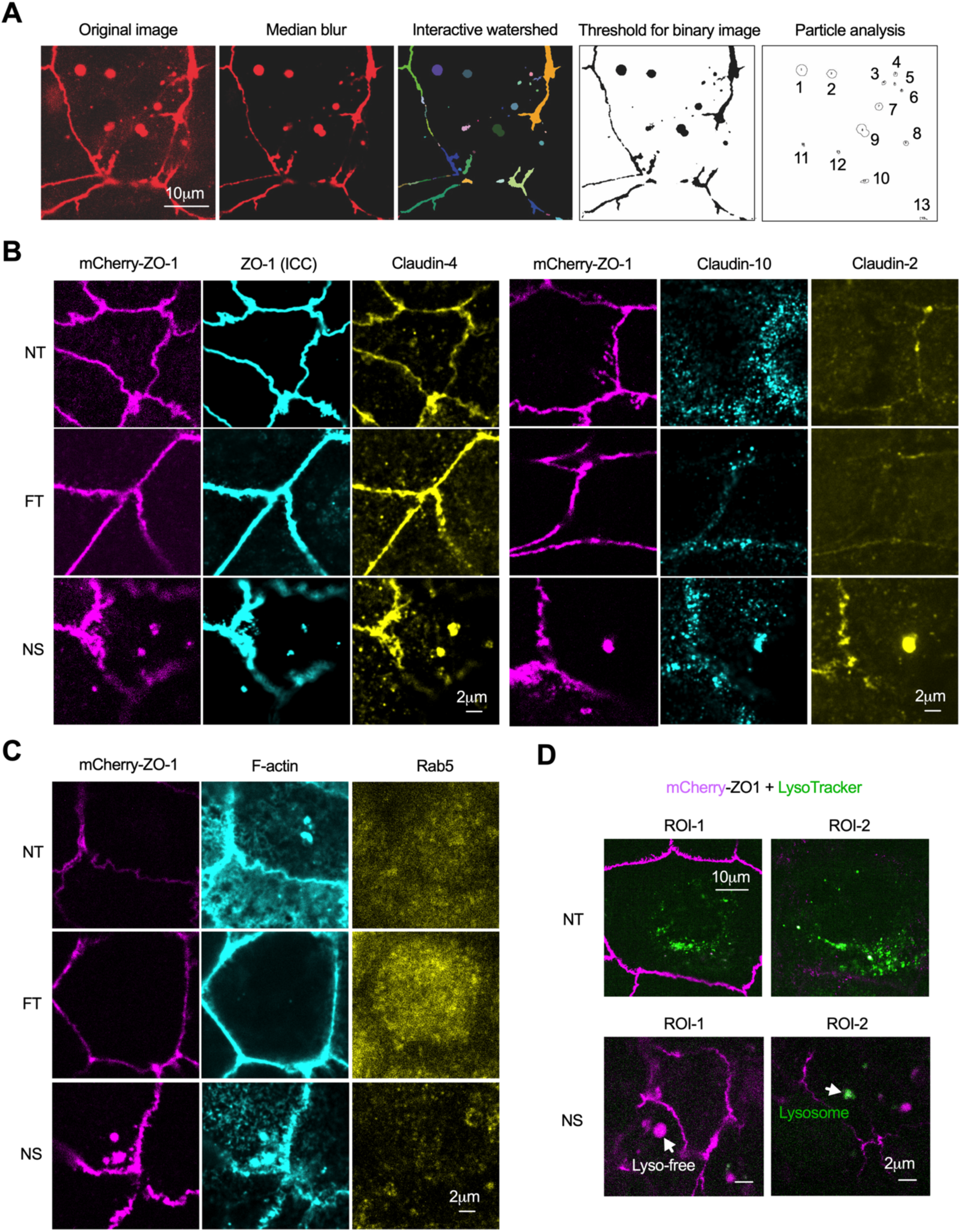
Quantification and compositional identification of cytosolic ZO-1 complexes formed upon nanostructured film treatment. (**A**) Processing and quantification of cytosolic complexes in images of engineered Caco-2 cell apical locations. Images were individually processed through median blur, interactive watershed, threshold for binary image and particle analysis. Size distribution of spherical complexes in images were reported in Fig. 2d,e. (**B,C**) Immunostained images of mCherry-ZO-1 engineered Caco-2 cells after treatment with NS film at 37°C for 1 hour showed co-localization of Claudin-2, Claudin-4, Claudin-10 (B) and F-actin (C) in cytosolic mCherry-ZO-1 complexes, while Rab5 was absent (C). (**D**) Images of LysoTracker dye stained live Caco-2 cells that were non-treated (NT) or in contact with NS film for 0.5-1.5 hour at 37°C showing that the apical cytosolic complexes are non-acidic. Scale bar: 2 μm.

**Fig. S4.**
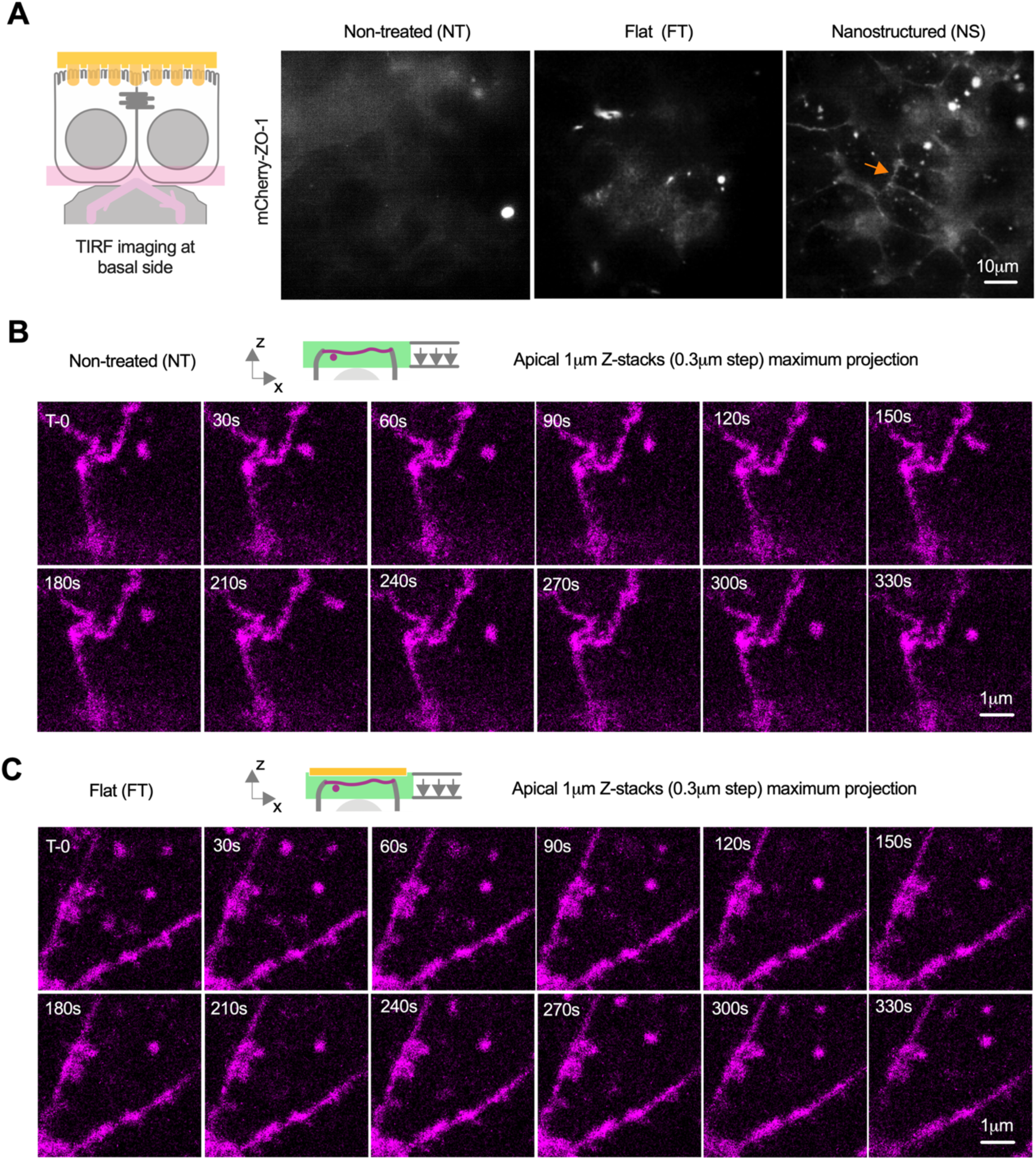
Live imaging of engineered mCherry-ZO1 cells at apical and basal side upon different treatment. (**A**) TIRF images at the basal side of engineered Caco-2 cells that were non-treated, or treated with flat or nanostructured film at the apical side for 1 hour at 37°C, showed interesting junction-like morphology (orange arrow) of mCherry-ZO-1 in cells stimulated by nanotopography. (**B,C**) Time lapse of apical mCherry-tagged ZO-1 in non-treated (NT) Caco-2 cells and those treated with flat film (FT) for 0.5-1.5 hour at 37°C. Fewer and smaller cytosolic structures of ZO-1 at apical side were observed in NT (**B**) or FT (**C**)-treated cells, and the interaction between tight junction and cytosolic structure was minimal. Images are projections of maximum pixel intensity of apical z-stack images acquired with 1 μm depth and 0.3 μm intervals.

**Fig. S5.**
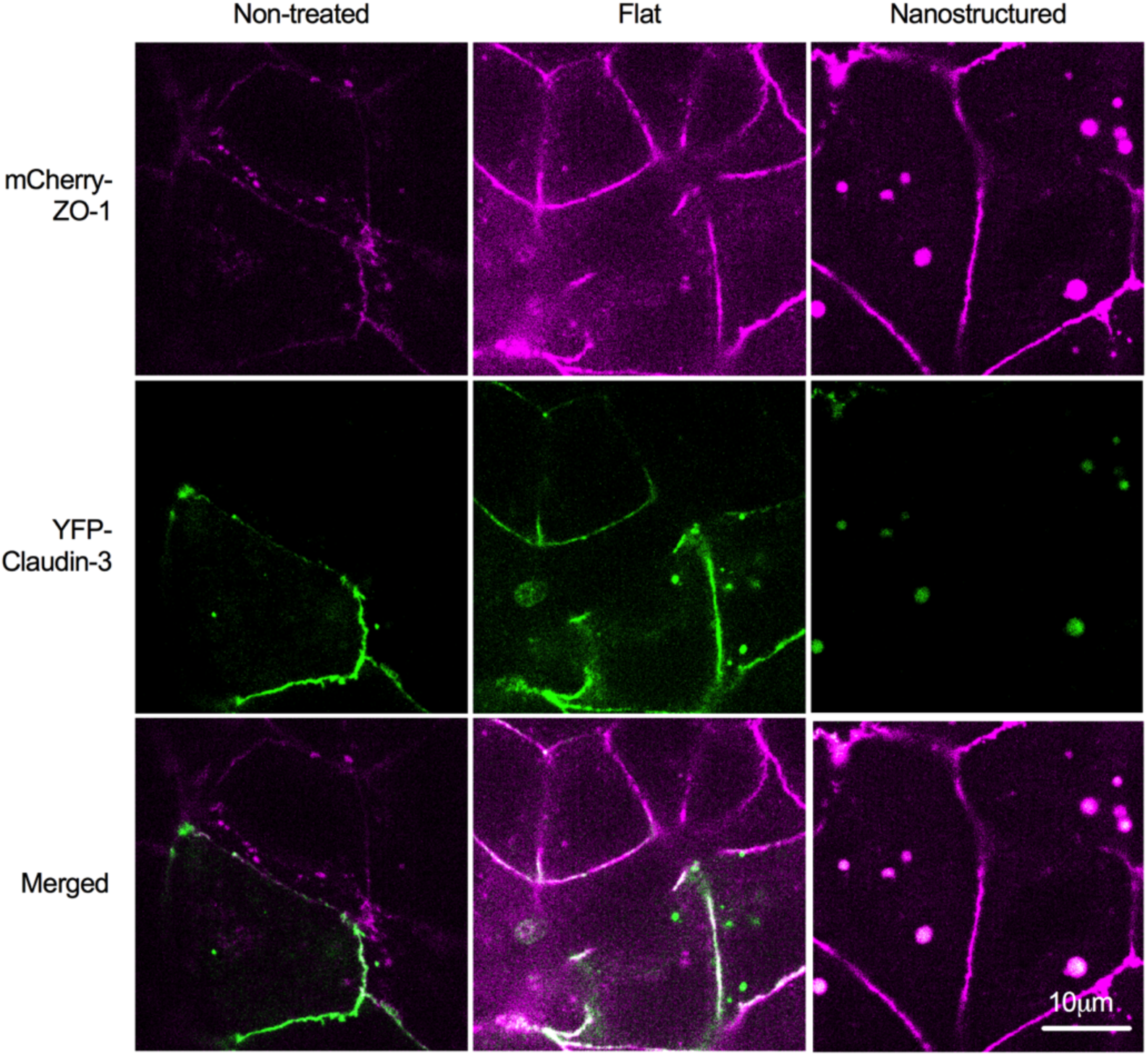
Live imaging of engineered mCherry-ZO1 cells that are exogenously transfected with YFP-Claudin-3, in response to flat and nanostructured film in comparison with untreated control. Adenovirus transfection of Caco-2 cells was controlled at 100 MOI (multiplicity of infection) to minimize unnatural accumulation of cytosolic Claudin-3, while tolerating uneven expression level among cells. Nanotopographic treatment led to apical cytosolic complexes that contain both ZO-1 and Claudin-3; however, in untreated or flat film treated cells, smaller and fewer cytosolic structures did not contain both proteins.

**Fig. S6.**
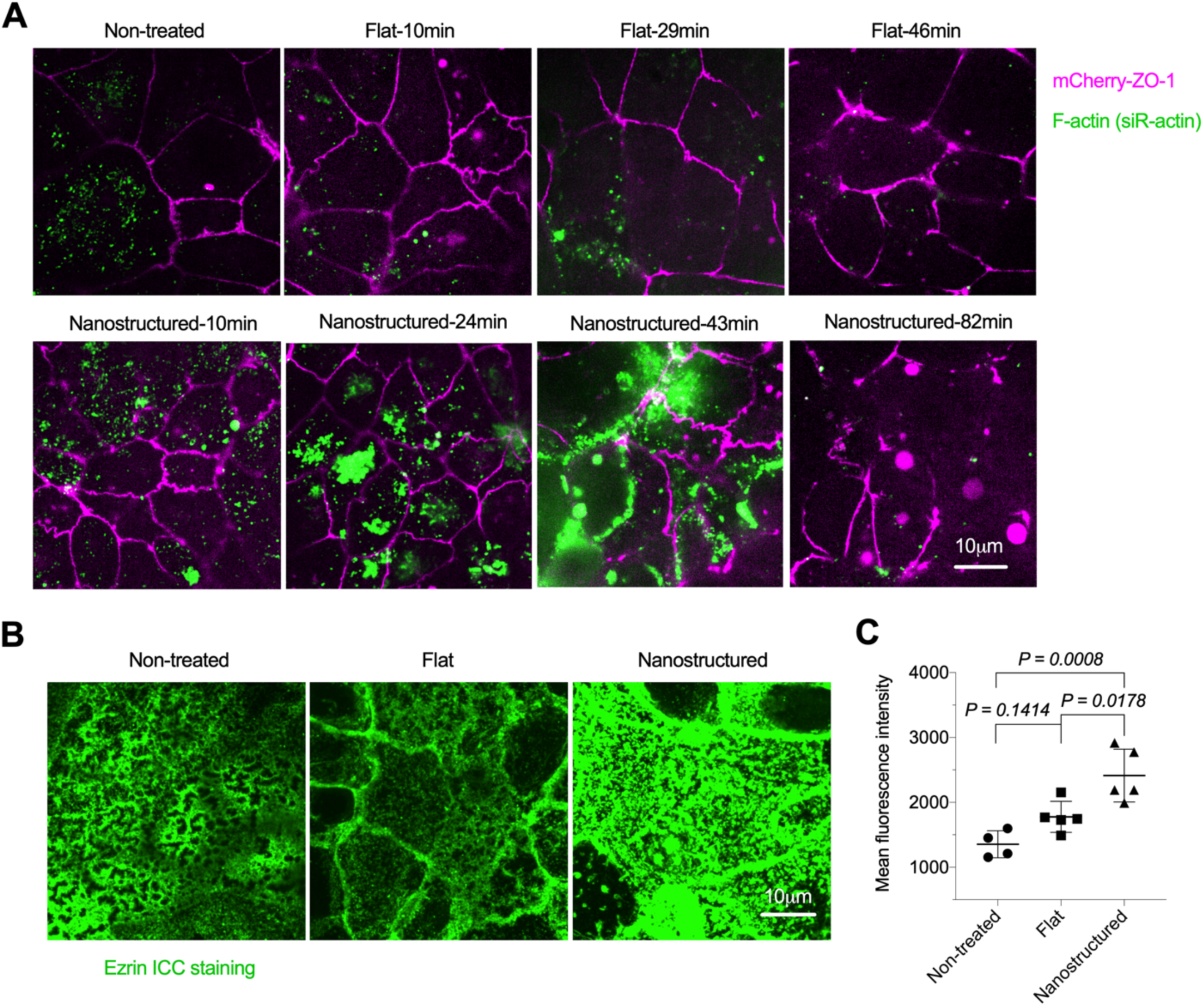
F-actin clusters and remodels at apical side upon nanostructured film treatment, while ezrin protein, a connecter between cell membrane protein and actin, was up-regulated as well. (**A**) Images of F-actin stained live mCherry-ZO-1 cells at different time point after flat and nanostructured film treatment. Apparent clustering and remodeling of F-actin in live cell apical side was observed from nanostructured film treatment; however, this trend was less obvious from flat film treatment and even less for non-treated cells. Images are projections of maximum pixel intensity of apical z-stack images acquired with 1 μm depth and 0.3 μm intervals. (**B**) Images of ezrin staining of Caco-2 cells treated with flat and nanostructured film for 1 hour at 37°C, compared with non-treated control. Images are projections of maximum pixel intensity of apical z-stack images acquired with 6 μm depth and 0.3 μm intervals. (**C**) Quantification of mean pixel intensity in images shown in (B). Data are mean ± s.d. (n = 5 images), and *P* values were determined by one-way ANOVA analysis.

### Note S1. Sequence of repair plasmid for HDR-based mCherry knock-in

The following 2747bp gene was cloned in pUC57-Kan plasmid by EcoRV (Genscript, Inc).

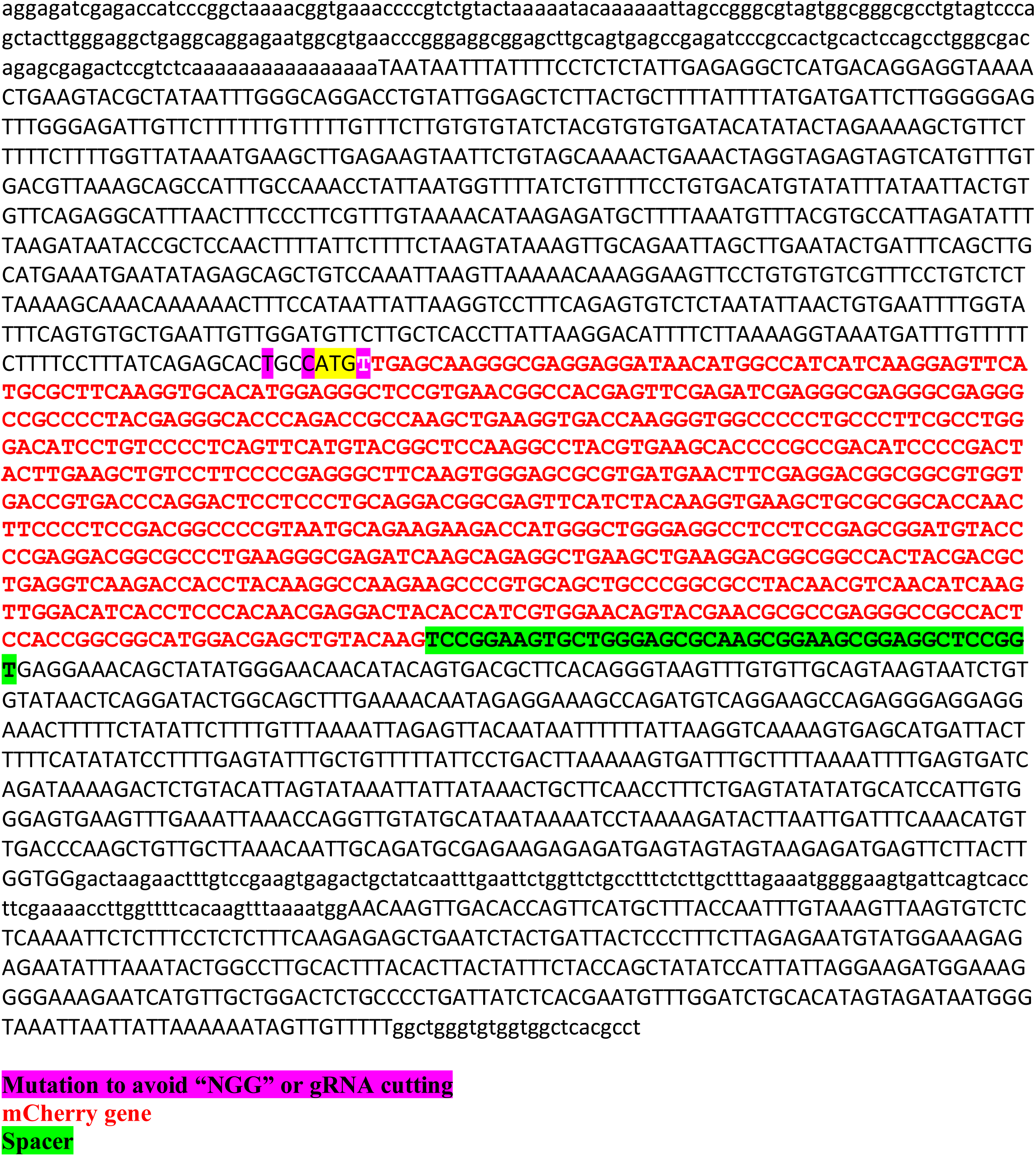

**Movie S1. Time-lapse movie of mCherry-tagged ZO-1 at the apical side of Caco-2 cells stimulated by nanotopographic cues, showing cytosolic ZO-1 proteins cluster into puncta within a network next to junctions. This movie represents images in Fig. 3A-ii.**

**Movie S2. Time-lapse movie of mCherry-tagged ZO-1 at the apical side of Caco-2 cells stimulated by nanotopographic cues, showing apical cytosolic complex circulates while collecting smaller ZO-1 structures which then moves basolaterally. This movie represents images in Fig. 3A-iv.**

**Movie S3. Time-lapse movie of mCherry-tagged ZO-1 at the apical side of Caco-2 cells stimulated by nanotopographic cues, showing active remodeling of cytosolic complexes and their interaction with junction-associated structures.**

**Movie S4. Time-lapse movie of mCherry-tagged ZO-1 (purple) and YFP-tagged claudin-3 (green) at the apical side of Caco-2 cells stimulated by nanotopographic cues, showing the involvement of both proteins in the initiation of complex formation and cytosolic migration. This movie represents images in Fig. 4C.**

**Movie S5. Time-lapse movie of mCherry-tagged ZO-1 (purple) and live-stained F-actin (green) at the apical side of Caco-2 cells stimulated by nanotopographic cues, showing: i) disappearance of ZO-1 concomitant with the appearance of F-actin; ii) F-actin wrapped around ZO-1 in cytosolic complex next to a tight junction; and iii) a large ZO-1 cytosolic complex lack of F-actin. This movie represents images in Fig. 5B.**

